# Detection of velogenic avian paramyxoviruses in rock doves in New York City, New York

**DOI:** 10.1101/2021.11.02.467002

**Authors:** Isabel Francisco, Shatoni Bailey, Teresa Bautista, Djenabou Diallo, Jesus Gonzalez, Joel Gonzalez, Ericka Kirkpatrick Roubidoux, Paul K. Ajayi, Randy A. Albrecht, Rita McMahon, Florian Krammer, Christine Marizzi

**Affiliations:** New York City Virus Hunters Program, BioBus, New York, NY 10027, USA; Department of Microbiology, Icahn School of Medicine at Mount Sinai, New York, NY 10029, USA; Central Park East High School, New York City Department of Education, New York, NY 10028, USA; High School for Environmental Studies, New York City Department of Education, New York, NY 10019, USA; Department of Infectious Diseases, St. Jude Children’s Research Hospital, Memphis, TN 38105, USA; The Global Health and Emerging Pathogens Institute, Icahn School of Medicine at Mount Sinai, New York, NY 10029, USA; Wild Bird Fund, New York, NY 10024, USA; Department of Pathology, Molecular and Cell Based Medicine Icahn School of Medicine at Mount Sinai, New York, NY 10029, USA

**Keywords:** birds, urban viral surveillance, avian paramyxovirus 1 (APMV-1), Newcastle Disease Virus (NDV), wildlife, community science, citizen science

## Abstract

Avian paramyxovirus 1 (APMV-1), also known as Newcastle disease virus (NDV), causes severe and economically important disease in poultry around the globe. Although a limited amount of APMV-1 strains in urban areas have been characterized, the role of the urban wild bird population as an APMV-1 reservoir is unclear. Since urban birds may have an important role for long-term circulation of the virus, fecal and swab samples were collected by community scientists from wild birds in New York City (NYC), New York, United States. These samples were screened for APMV-1 and genotypically characterized by sequencing of the complete genome. A total of 885 samples were collected from NYC parks and from a local wildlife rehabilitation clinic from October 2020 through June 2021. Eight birds (1.1 %) screened positive for the APMV-1 nucleoprotein gene by conventional reverse transcription polymerase chain reaction (RT-PCR), and two live viruses were isolated via egg culture. The F protein cleavage site, an indicator of pathogenicity, was present in the two samples fully sequenced by next generation sequencing, and positioned ^112^R R K K R F^117^. Phylogenetic analysis of the F gene coding sequence classified both isolates into genotype VI, a diverse and predominant genotype responsible for NDV outbreaks in pigeon and dove species worldwide.

**Importance:** Here we describe the first large-scale effort to screen for APMV-1 in New York City’s wild bird population as part of the New York City Virus Hunters program, a community science initiative. We have characterized two isolates of NDV, with phylogenetic analyses suggesting diversity in established and circulating strains of pigeon paramyxoviruses. Our isolates are also domestic reference strains for future APMV-1 vaccine developments. Future surveillance in this region may contribute to our understanding of NDV’s evolution and genetic diversity, as well as inform poultry husbandry and vaccination practices in New York State.

## Introduction

Avian Paramyxovirus 1 (APMV-1), also known as Newcastle Disease Virus (NDV), is an economically important poultry pathogen, with occasional outbreaks reported in wild birds (1, 2). As a causative agent of virulent Newcastle disease (ND), it often causes neurological symptoms in birds, including but not limited to twisting of the head and neck, poor balance, tremors, and paralysis of wings and legs (2). Although NDV can be controlled through vaccination and maintaining strict biosecurity measures, it still poses a high economic burden on the poultry industry (3–5). Chickens have been reported as highly susceptible, as well as gallinaceous birds such as turkey, quail, and guinea. The 2003 NDV outbreak in the western United States alone resulted in the death or culling of over 3 million birds (6–9). Distributed around the globe, NDV is an Office International des Epizooties (OIE) notifiable disease and prompt reporting of any outbreak is mandatory to local regulatory agencies (9, 10). In the United States, a sporadic form of the disease exists throughout the year, and only a limited number of outbreaks are officially reported annually to the United States Department of Agriculture (USDA) (9). Most recent outbreaks confirmed by the USDA include more than 470 premises in California, including 4 commercial premises in 2020 (9).

APMV-1 is a member of the genus *Avulavirus* within the family *Paramyxoviridae*, order *Mononegavirales* (11). The enveloped genome of this single-stranded negative sense RNA virus is approximately 15.2 kb in length and encodes for 6 different proteins, i.e. nucleocapsid protein (NP), phosphoprotein (P), fusion protein (F), matrix protein (M), hemagglutinin-neuraminidase (HN), and the RNA polymerase (L) protein (11). The NP protein forms the nucleocapsid core with genomic RNA, to which P and L proteins are bound (12). The non-glycosylated M protein is located beneath the envelope and associated with virus assembly and budding (12). The two surface glycoproteins HN and F are responsible for binding to host cell sialic acid receptors (HN) and for fusion of the viral envelope to the host cell membrane (F) (13). As a property of the family, APMV-1 carries high protein coding capacity, which is further enhanced by RNA editing, resulting in generation of the two non-structural proteins V and W during the transcription of the P gene (13, 14). While the V protein is a key regulator of cell apoptosis and viral replication (14), very little is known about the W protein (15).

Several NDV molecular classification systems have been developed in order to document and track this virus’s genetic diversity and evolution. Unified phylogenetic classification criteria were established by the OIE, separating all existing NDV isolates into two classes (class I and class II) and as many as twenty-one genotypes, as described by Dimitrov et al (16). This collaborative effort also suggests updated guidelines for nomenclature, especially for sub-genotypes, as the worldwide circulation and evolution of NDV will continue to lead to the emergence of new NDV variants.

On the basis of conventional *in vivo* pathogenicity indices for poultry, APMV-1 strains are classified into several pathotypes. Viscerotropic velogenic APMV-1 is highly pathogenic and causes intestinal infection with high mortality in birds, whereas neurotropic velogenic APMV-1 is responsible for symptoms of the respiratory and nervous systems with high mortality. The mesogenic strains are less pathogenic, often with acute respiratory and nervous symptoms but with relatively low mortality. The lentogenic strains of APMV-1 cause mild respiratory tract infections, allowing for a prolonged virus replication and shedding (9, 10). This wide range in pathogenicity has been attributed to differences in the F protein cleavage site (10). While all mesogenic and velogenic APMV-1 strains carry an amino acid sequence of ^112^R/K-R-Q-R/K-R-F^117^ within the F protein, lentogenic strains are characterized by ^112^G/E-K/R-Q-G/E-R-L^117^ (10).

While no treatment for NDV infection is known, it can be controlled by the use of vaccines, and several prophylactic NDV vaccines are available on the national and international market for use in commercial poultry (10). Live attenuated NDV vaccines are suitable for mass production and utilized for routine mass vaccination via spray or drinking water. [17].Despite the extensive and unrestricted use of vaccines to prevent NDV in domestic poultry in the United States, APMV-1 still remains one of the main poultry diseases in both commercial and backyard chickens (9). This might be explained by the mismatch between field and vaccine strains and emergence of novel APMV-1 strains by divergence of sub-genotypes circulating in vaccinated poultry. Dimitrov *et al*. reported in 2016 that vaccine strains currently used are three to seven decades old, and up to 26.6% genetically distant (nucleotide distance) from virulent contemporary NDV strains (3, 17).

Moreover, similar to rural wild birds, the role of wild urban birds in the epizootiology of APMV-1 has remained understudied and therefore unclear. Wild birds represent major natural reservoirs and are potential dispersers of infectious disease, including pathogens like APMV-1 and avian influenza virus. Densely populated urban areas and city parks are unique habitats for wild birds, where species in the order of *Anseriformes* (e.g. ducks, geese and swans) and *Charadriiformes* (e.g. shorebirds and gulls) are more likely to come into contact with other bird species. Virulent strains of APMV-1 are often maintained in wild birds in close proximity to water such as cormorants and gulls. Strains from class II genotype VI of APMV-1, sometimes called pigeon paramyxovirus 1 (PPMV-1), are also believed to be endemic in locations with large populations of *Columbiformes* or pigeons and doves (18), species ubiquitous to New York City. To date, there has been no spillover outbreak from *Columbiformes* species into poultry documented in the US, though the role of wild birds in the transmission of virulent NDV is not fully understood (19, 20).

In North America, millions of wild birds migrate along one of the four north-south flyways annually - the Atlantic, Mississippi, Central, and Pacific (21). Along the Atlantic flyway, there are many key sites that migratory birds utilize to gather to breed, feed, or rest. With major metropolitan areas such as Massachusetts and New York along the route, it is also the flyway most densely populated by humans. Migratory birds resting in urban areas may be particularly important in the transmission pathway among immunologically naïve birds, because once infected, they may shed virus particles for weeks through fecal and respiratory droppings, and without showing any clinical symptoms. No avian surveillance has established what viruses circulate endemically in the migratory species located in New York City.

Here, we report results from the first large-scale surveillance investigation on APMV-1 in wild birds performed in the New York City metropolitan area. Launched in 2020, New York City Virus Hunters is a community science program that is based on community participation. A collaboration between BioBus, a Science Outreach organization; the Krammer Laboratory at the Icahn School of Medicine at Mount Sinai, an influenza research laboratory; and the Wild Bird Fund (WBF), an urban wildlife rehabilitation clinic, this study aims to address the lack of extensive baseline data for avian viruses in wild birds in urban areas. In order to evaluate the degree of genetic diversity of APMV-1 strains circulating in NYC’s wild bird populations and to estimate the relationships to APMV-1 strains that circulated in the Northeast region in the past, the complete genomes of two APMVs isolated from NYC birds were characterized phylogenetically. Finally, this program also addresses the lack of participatory research opportunities for the local community to help to prepare for and prevent the next pandemic.

Participatory research, sometimes termed citizen science or community science, has become an important data source in many scientific disciplines (22, 23). In contrast to other projects where participants partake in only data collection, the New York City Virus Hunters program invited participants to be actively involved in every step of the research process. New York City residents engaged in study design and outreach strategies, followed by trained and supervised sample collection, processing and analysis, as well as data dissemination tailored to the scientific community and the general public. This closed-feedback loop ensures that program participants take the data they generate directly to their communities, potentially improving awareness about infectious disease, pandemic preparedness and in long-term vaccination rates among the least-vaccinated and therefore vulnerable populations.

## Materials and Methods

### Study sites and sample collection

New York City Parks and natural areas were sampled for wild bird fecal samples from October 2020 through June 2021, including seven parks in Manhattan, Brooklyn, and the Bronx **(Figure 1)**. Visited parks were Central Park, Fort Tryon Park, George Washington Park and Carl Schurz Park in Manhattan; Van Cortlandt Park in the Bronx; and Prospect Park in Brooklyn. Fecal samples were collected using sterilized microcentrifuge tubes and cotton swabs. Samples were taken at least 10 feet apart and fresh or visibly moist samples were preferentially selected. Fecal samples were transported on ice and kept at −80 °C until processing.

**Figure 1:**
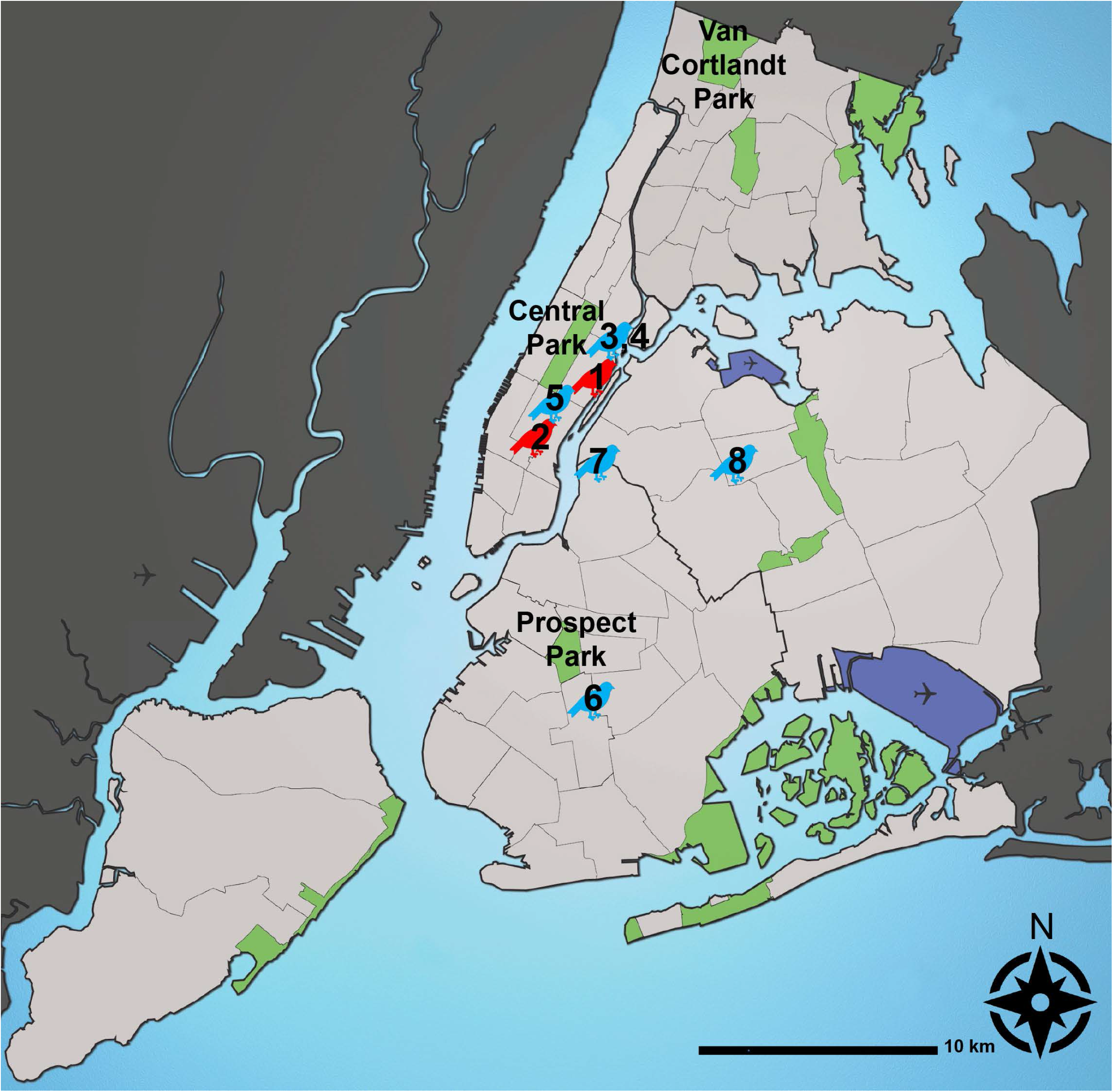
Sampling location of birds that confirmed positive for APMV in New York City by RT-PCR targeting the fusion (F) gene. Six samples that screened positive for APMV are indicated in blue. Two samples with additionally available full length genomic APMV sequences are indicated in red. Two samples where no further location than New York City was available (samples 3 and 4), are mapped to the location of the intake site (Wild Bird Fund). 1: 21-0109: East 71st Street, Manhattan, New York, NY 2: 21-0052: West 33rd and Broadway, Manhattan, New York, NY 3,4: 21-0490 and 21-0550: Manhattan, New York, NY 5: 21-0705: West 58^th^ Street and 6^th^ Avenue, Manhattan, New York, NY 6: 21-0693: Newkirk Plaza, Flatbush, Brooklyn, NY 7: 21-0710: Clay Street, Greenpoint, Brooklyn, NY 8: 21-0593: Van Kleeck Street, Elmhurst, Queens, NY Airports: North East: LGA South East: JFK West: EWR, major parks and natural areas are indicated in green, top sampling location for each borough labeled (Central Park in Manhattan, NY; Prospect Park in Brooklyn, NY and Van Cortlandt Park in the Bronx, NY). Location approximate. Bird icon not to scale. Graph prepared by Christine Marizzi, based on https://commons.wikimedia.org/wiki/File:Waterways_New_York_City_Map_Julius_Schorzman.png; CC-BY-SA-2.5

Wild birds surrendered to the WBF in Manhattan, NY for rehabilitation were opportunistically sampled from January through June 2021. Fresh fecal samples (≤ 12 hours) were collected from individual clinic enclosures to sample live birds in rehabilitation. Cloacal and oropharyngeal swabs were also collected from recently deceased or euthanized birds using sterile flocked nylon-tipped swabs. Swab tips were placed in conical tubes containing either MicroTest viral transport medium (Thermo Scientific, USA) or medium containing 50% phosphate buffered saline and 50% glycerol, supplemented with 1% Gibco™ Antibiotic-Antimycotic 100X (Thermo Scientific, USA)). Fecal samples and swab tips were kept in a 4 °C freezer for pickup and then transferred to −80 °C within 48 hours of collection. All live bird sampling was performed or supervised by New York State licensed wildlife rehabilitators employed by the WBF.

### RNA extraction and RT-PCR

Fecal samples were diluted in phosphate-buffered saline, pH 7.4 (1X, Thermo Scientific, USA) for processing. Suspended fecal samples and swab samples were centrifuged at 4,000 x g for 15 min and viral RNA was extracted from each supernatant using the QIAamp Viral RNA Mini Kit (Qiagen, USA). Conventional RT-PCR was performed using the Invitrogen™ SuperScript™ IV First-Strand Synthesis System (Thermo Scientific, USA) for cDNA synthesis and DreamTaq Green PCR Master Mix (2X) (Thermo Scientific, USA) for RT-PCR, using previously described primers for APMV-1 surveillance that target the nucleoprotein (NP) gene **(Table 1)**. cDNA was synthesized at 55 °C for 10 min. Cycling conditions for APMV-1 PCR consisted of a pre-denaturation step at 95 °C for 1 min, followed by 40 cycles of denaturation at 95 °C for 1 min, annealing at 50 °C for 30 sec, and extension at 72 °C for 30 sec, with a final extension step at 72 °C for 5 min. PCR amplicons were visualized with SYBR Safe™ DNA Gel Stain in 2% Ultra Pure Agarose (Thermo Scientific, USA). DNA bands were excised and purified using the QIAquick Gel Extraction Kit (Qiagen, USA) and sent for commercial Sanger Sequencing through Genewiz^®^ to verify the identity of samples that screened positive. Samples were also screened for avian influenza virus, with results appearing elsewhere.

**Table 1.**
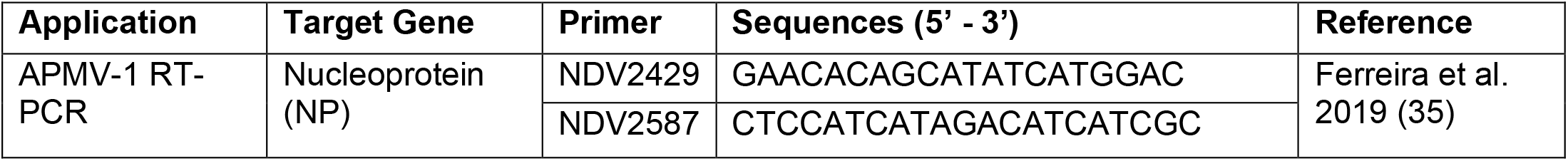
Primer set used for APMV-1 virus detection by RT-PCR.

### Signal amplification and sequencing

Supernatants for samples that screened positive on PCR were inoculated into the allantoic cavity of 10-11-day-old embryonated specific-pathogen-free chicken eggs and incubated at 37 °C for 48 hours to amplify the signal detected in the initial samples. Allantoic fluid from incubated eggs was harvested and centrifuged at 4,000 x g for 15 min. Presence of virus was then determined by a hemagglutination assay using chicken red blood cells and standard methods. Viral RNA was extracted from virus-infected allantoic fluid and conventional RT-PCR was used to confirm the presence of APMV-1 as described above.

For isolated APMV-1, primer sets were designed for this study in order to obtain the full coding sequence of the fusion (F) gene for APMV-1. Primers were synthesized by Thermo Scientific, USA. Conventional RT-PCR and commercial Sanger sequencing were performed using these primer sets as described above. All infectious materials were maintained in Biosafety Level 2+ containment at the Icahn School of Medicine, Mount Sinai. Any sample with an F-gene containing a polybasic cleavage site (indicative of non-lentogenic NDV) was immediately reported to the USDA and transferred to a BSL3+ select agent facility for storage.

RNA from two confirmed APMV-1 cases was also submitted for Next Generation Sequencing (NGS) through Genewiz^®^. Sequence reads were trimmed to remove possible adapter sequences and nucleotides with poor quality using Trimmomatic v.0.36. The trimmed reads generated were aligned in STAR aligner v.2.5.2b to a closely related North American pigeon paramyxovirus reference genome in GenBank with accession number KP780874.2. Consensus sequences were generated from the alignment files in the software Geneious V.2021.1.1 (Biomatters, Inc., New Zealand). These consensus sequences were also verified using V-pipe, a bioinformatics pipeline designed for the analysis of NGS data from RNA viruses (24).

### Phylogenetic analysis

F gene coding sequences for the two APMV-1 isolates were compared with the online NIH GenBank database using BLASTN. Genotype and sub-genotype classification were performed according to criteria for the updated unified phylogenetic classification for NDV as described by Dimitrov *et al* (16). Alignments were performed using the ClustalW program and visualized in MEGA X. Phylogenetic trees were created and visualized in MEGA X using the Maximum Likelihood (ML) method and the Tamura-Nei model. Evolutionary distances between groups were inferred utilizing the Maximum Composite Likelihood model with rate variation among sites modeled with a gamma distribution (shape parameter = 1). Full genome sequence alignments with a representative reference genome were performed with Clustal Omega at EMBL-EBI using default settings (25–27).

## Results

### Sampling and virus detection

A total of 885 birds were sampled from October 2020 to June 2021. 74 fecal samples from three NYC Parks, 65 fecal samples from the Wild Bird Fund, and 116 swab samples (58 oropharyngeal and 58 cloacal) from the WBF were screened for APMV-1 by RT-PCR (N=255) (**Figure 1**). All WBF samples were from birds rescued in NYC and admitted to the site, which is located in Manhattan (**Figure 1**). 36 of the 74 fecal samples collected in parks (48.6%) were from Canada geese (*Bratana canadensis*), while 136 of the 197 birds sampled at the WBF (69.0%) were rock doves (*Columbia livia*).

Of the 255 samples processed, the avian paramyxovirus-1 nucleoprotein (NP) gene was detected in swab samples from eight birds (4.1%) spanning three NYC boroughs (**Table 2**). Seven of the APMV-1-positive birds were pigeon species and one was an American woodcock (*Scolopax minor*, **Table 2**). Of the samples positive on RT-PCR, two APMV-1 viruses were isolated via chicken egg inoculation, both positive on HA assay after the first egg passage.

**Table 2.**
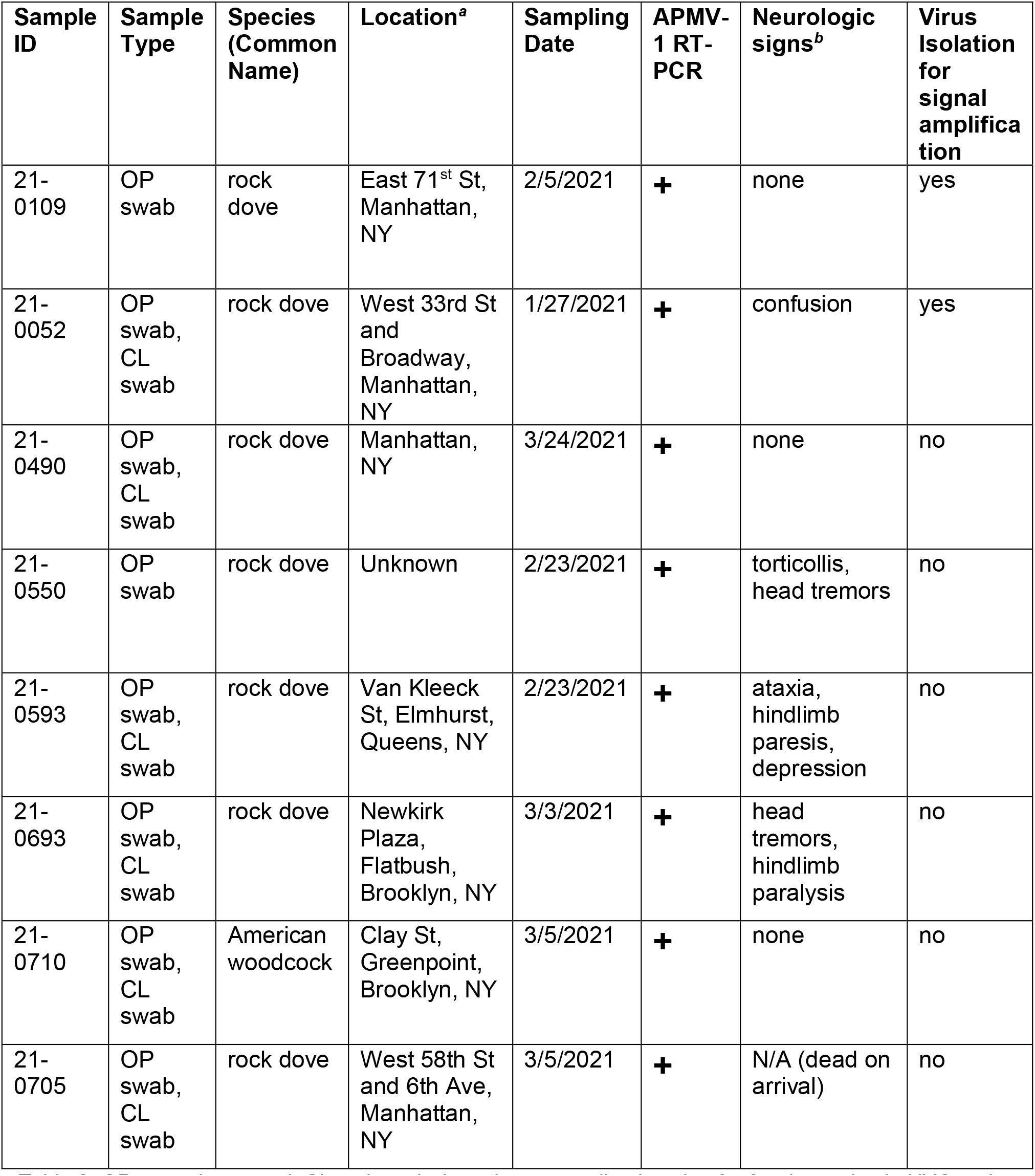
Clinical and sampling information for wild birds positive on RT-PCR for avian paramyxovirus-1 (APMV-1) in New York City from October 2020 to June 2021.

| Sample ID | Sample Type | Species (Common Name) | Location<sup>a</sup> | Sampling Date | APMV-1 RT-PCR | Neurologic signs<sup>b</sup> | Virus Isolation for signal amplification |
| --- | --- | --- | --- | --- | --- | --- | --- |
| 21-0109 | OP swab | rock dove | East 71<sup>st</sup> St, Manhattan, NY | 2/5/2021 | + | none | yes |
| 21-0052 | OP swab, CL swab | rock dove | West 33rd St and Broadway, Manhattan, NY | 1/27/2021 | + | confusion | yes |
| 21-0490 | OP swab, CL swab | rock dove | Manhattan, NY | 3/24/2021 | + | none | no |
| 21-0550 | OP swab | rock dove | Unknown | 2/23/2021 | + | torticollis, head tremors | no |
| 21-0593 | OP swab, CL swab | rock dove | Van Kleeck St, Elmhurst, Queens, NY | 2/23/2021 | + | ataxia, hindlimb paresis, depression | no |
| 21-0693 | OP swab, CL swab | rock dove | Newkirk Plaza, Flatbush, Brooklyn, NY | 3/3/2021 | + | head tremors, hindlimb paralysis | no |
| 21-0710 | OP swab, CL swab | American woodcock | Clay St, Greenpoint, Brooklyn, NY | 3/5/2021 | + | none | no |
| 21-0705 | OP swab, CL swab | rock dove | West 58th St and 6th Ave, Manhattan, NY | 3/5/2021 | + | N/A (dead on arrival) | no |

### Sequencing

The full F gene coding sequences for both isolated APMV-1 viruses were obtained via Sanger sequencing. Both sequences are 1659 bp in length and encode a predicted F protein of 553 amino acids. Both isolates also presented fusion protein cleavage sites compatible with virulent Newcastle Disease virus (NDV), with five basic amino acids at positions 112-116 and a phenylalanine residue at position 117 (^112^R R K K R F^117^) (10). Both isolates were reported to the USDA as *APMV-1/Rock_Dove/NYC/USA/NYCVH/21-0109/2021* and *APMV-1/Rock_Dove/NYC/USA/NYCVH/21-0052/2021*. In accordance with federal regulations, the detection of virulent NDV was reported to the USDA and all raw sample materials for both birds were moved to a select agent biosafety level 3 facility within the Icahn School of Medicine at Mount Sinai.

The complete genome sequences of both APMV-1 isolates were obtained by commercial Next Generation Sequencing (NGS) from extracted RNA (≥100.000 fold coverage). Both isolates had an expected consensus genome sequence length of 15,192 nucleotides. Comparison of the F gene sequences for both viruses with the sequences previously obtained via RT-PCR confirmed the presence of a virulent cleavage site at the F protein **(Figure 2)**. The F protein coding sequence for one isolate, 21-0052, contained one difference in nucleotide, at position 5822, when compared with the complete genome obtained by NGS sequencing. The nucleotide and amino acid in question as indicated by the NGS data were identical to our reference sequences. The NGS sequence was used for further analysis of this isolate as well as the other isolate (21-0109), which was 100% identical to its Sanger sequence.

**Figure 2.**
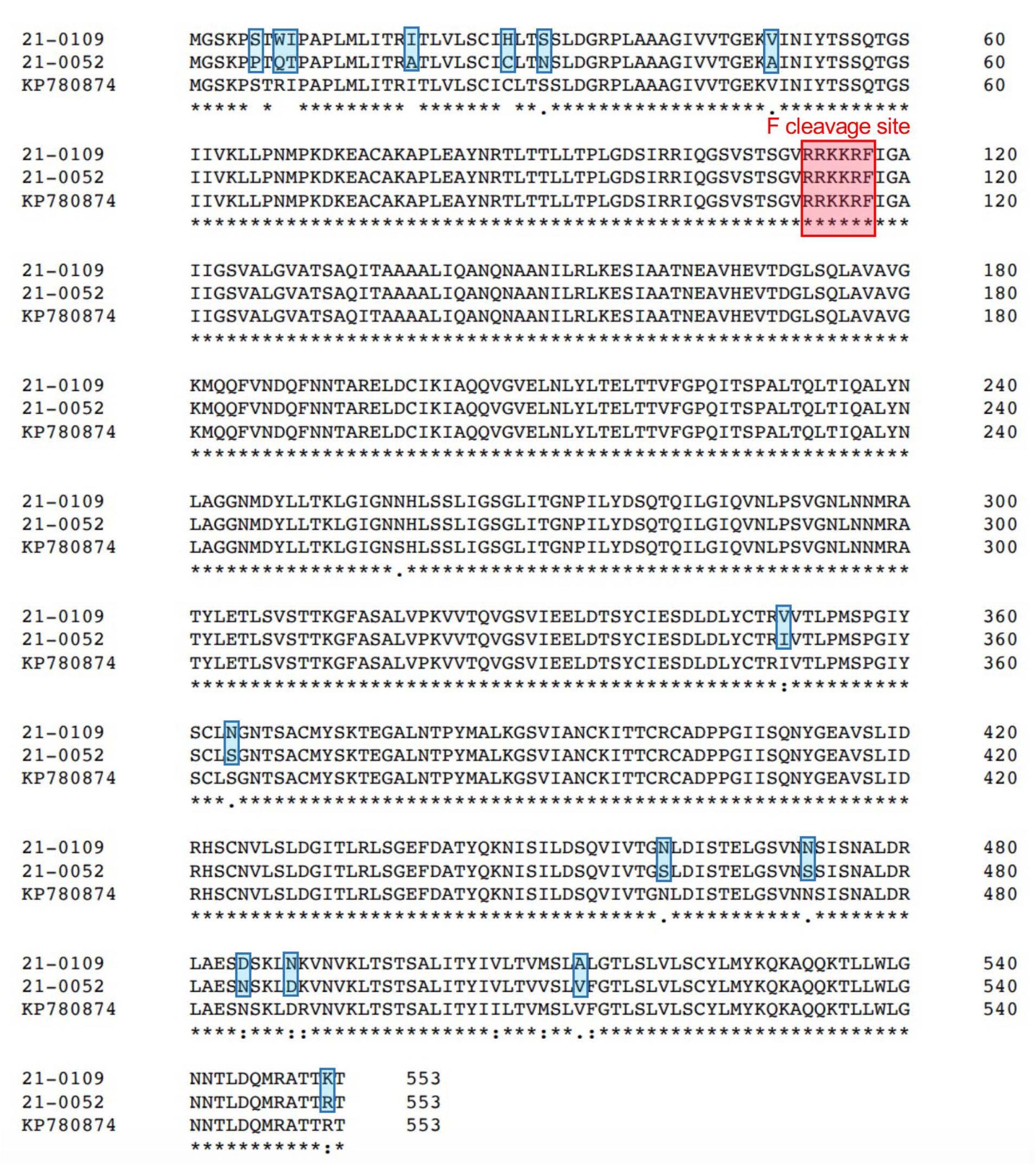
Characterization of F protein virulence determination site of isolates by comparison to a representative sequence of APMV-1. Single amino acid polymorphisms between the two novel APMV-1 isolates identified in this study are highlighted in blue. The typical RRKRR polybasic cleavage site motif is highlighted (red rectangle). Alignment was prepared with Clustal Omega at EMBL-EBI using default settings.

### Clinical features

45 of the 123 (36.6%) birds sampled at the WBF were documented to have neurologic symptoms while at the clinic, with a likely diagnosis (i.e. lead toxicity, head trauma) for 12 of these cases (26.7%). Of the eight birds that screened positive on RT-PCR for APMV-1, four (50%) presented with neurologic clinical signs without a likely diagnosis, including head tremors, torticollis, ataxia, paresis and paralysis (**Table 2**). Of the two birds with live APMV-1 isolated, one (21-0052) presented with generalized weakness and confusion; the other (21-0109) had no neurologic signs. Both birds were rock doves that presented to the WBF in January 2021 for generalized weakness and inability to fly, declined over time, and died in early February 2021.

### Phylogenetic analysis

Phylogenetic analysis of the two F gene coding sequences indicated that they belong to Class II, genotype VI, subgenotype VI.2.1.1.1, using the representative dataset for Class II NDV from the recently updated and unified NDV classification system (**Figure 3**) (16). The pairwise nucleotide distance between the F gene coding sequences for the two isolates was calculated to be 2.4%, meeting the criteria for the same subgenotype (≤5% difference).

**Figure 3.**
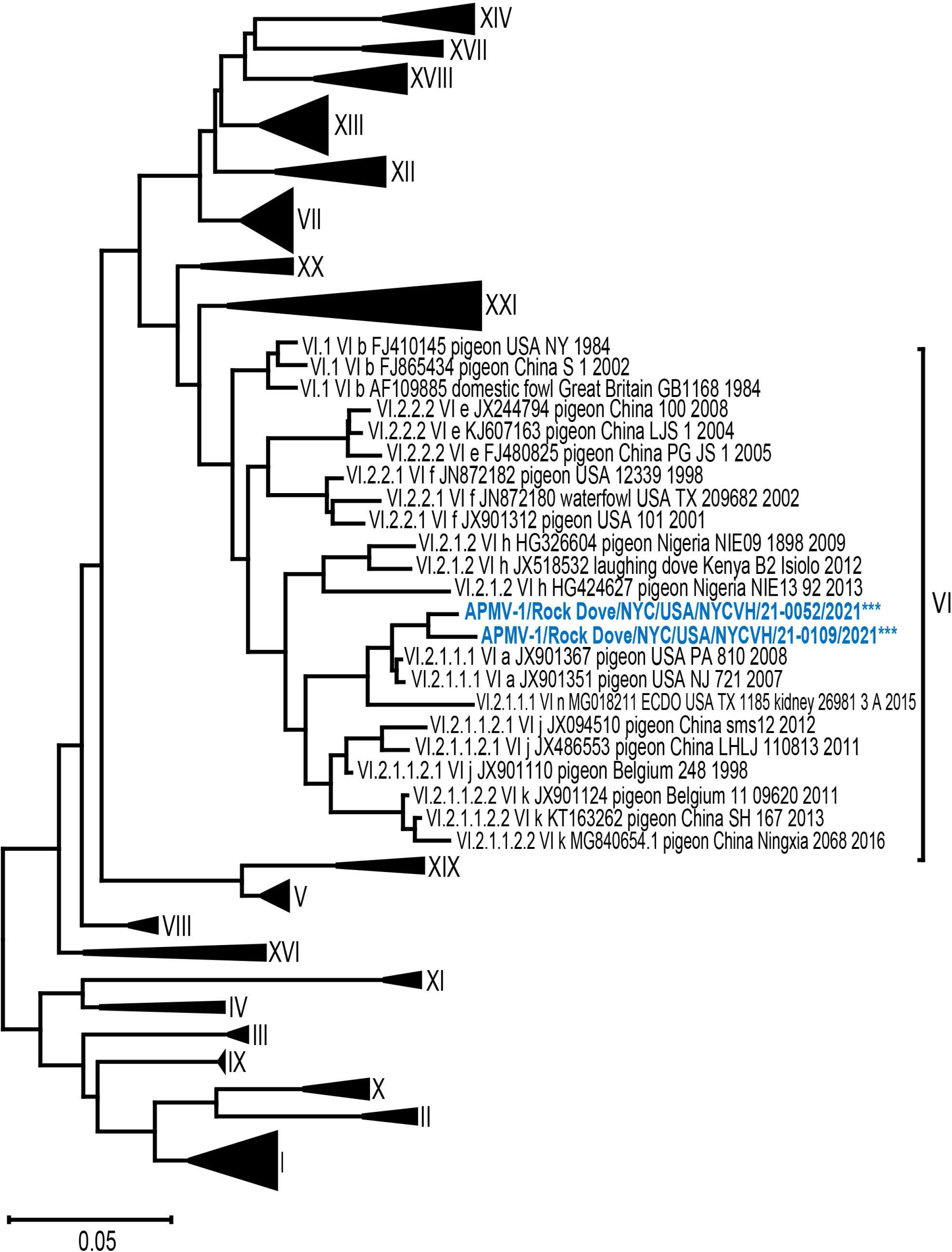
Phylogenetic tree of isolates with representative Class II NDV F protein sequences. Evolutionary analysis by Maximum Likelihood method and Tamura-Nei model. Tree includes the two APMV-1 isolates from this study and 125 representative F gene coding sequences from Class II NDV from the pilot dataset for genotype classification by Dimitrov *et al* (16) (N=127). The scale bar represents the percent divergence or nucleotide difference between sequences. The tree with the highest log likelihood (−31439.35) is shown. Initial tree(s) for the heuristic search were obtained automatically by applying Neighbor-Join and BioNJ algorithms to a matrix of pairwise distances estimated using the Tamura-Nei model, and then selecting the topology with superior log likelihood value. The tree is drawn to scale, with branch lengths measured in the number of substitutions per site. This analysis involved 127 nucleotide sequences. Codon positions included were 1st+2nd+3rd+Noncoding. There were a total of 1662 positions in the final dataset. Evolutionary analyses were conducted in MEGA X.

Nucleotide BLAST analysis of the F gene coding sequences for 21-0109 and 21-0052 showed close similarity to other NDV strains isolated from pigeons in the US in GenBank. Sixty-five F gene coding sequences were selected for further phylogenetic analysis, excluding sequences from earlier than 2007 (**Figure 4**). The evolutionary distances between our isolates, these selected sequences, and other genotype VI viruses in the consortium dataset were estimated and are included in table form in the supplemental materials (**Supplemental table S1**) (N=76). The average nucleotide distance between these sequences was 3.5%.

**Figure 4.**
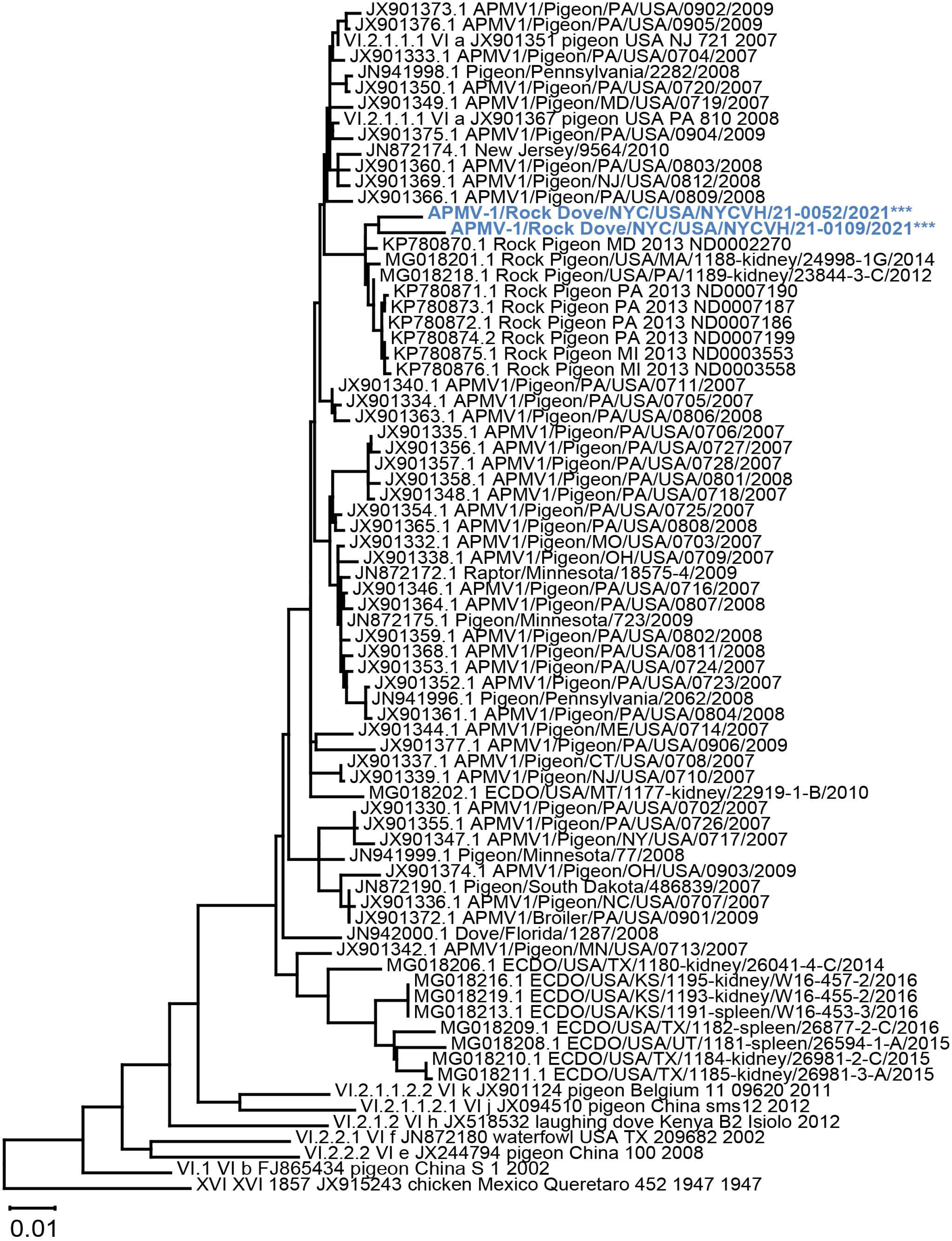
Phylogenetic tree of isolates with closely related NDV F protein sequences. Evolutionary analysis by Maximum Likelihood method and Tamura-Nei model. Tree includes F gene coding sequences from the two APMV-1 isolates from this study, 9 representative Class II Genotype VI NDV sequences from the pilot dataset by Dimitrov *et al*. (16), and 65 closely related NDV strains from wild birds in the USA. The scale bar represents the percent divergence or nucleotide difference between sequences. The tree with the highest log likelihood (−8126.65) is shown. Initial tree(s) for the heuristic search were obtained automatically by applying Neighbor-Join and BioNJ algorithms to a matrix of pairwise distances estimated using the Tamura-Nei model, and then selecting the topology with superior log likelihood value. The tree is drawn to scale, with branch lengths measured in the number of substitutions per site. This analysis involved 76 nucleotide sequences. Codon positions included were 1st+2nd+3rd+Noncoding. Positions containing gaps and missing data were eliminated. There were a total of 1656 positions in the final dataset. Evolutionary analyses were conducted in MEGA X.

Based on F gene coding sequences, the two isolates from this current study were most closely related to sub-genotype VI.2.1.1.1 sequences from Maryland (MD), Pennsylvania (PA) and Michigan (MI) from 2013, as well as sequences from Massachusetts (MA) and PA from 2014 and 2012, respectively. They were therefore more closely related to PPMV’s from other states than to each other. The two isolates expectedly grouped together with these closely related sequences (**Figure 4**). Our isolates also grouped together with NDV sequences from PA, New Jersey (NJ), MD, Minnesota (MN) and Missouri (MO) that were in the same subgenotype, while NDV sequences from Kansas (KS), Texas (TX), and Utah (UT) were grouped together in a separate subgenotype (**Figure 4**).

### Data Availability

The obtained complete genome sequences of *APMV-1/Rock_Dove/NYC/USA/NYCVH/21-0109/2021* and *APMV-1/Rock_Dove/NYC/USA/NYCVH/21-0052/2021* were submitted to NCBI NIH GenBank and are available under accession numbers XXXX to XXXX.

## Discussion and Outlook

The present study demonstrates the role of community science as a sentinel for urban viral surveillance initiatives that potentially could detect emerging infectious disease. A similar strategy of conducting surveillance on wild birds in urban spaces would be useful for other regions located along major flyways. Monitoring of those viruses that have the potential to infect a wide range of birds, and in some cases humans, could serve as an early warning system. Because local residents are engaged from the beginning of the project to the communication of scientific findings, this model also has potential to raise scientific literacy among community members, in particular increasing their understanding of infectious disease and environmental health information specific to an urban area.

We found that none of our fecal samples screened positive for APMV-1. Fecal samples for this study were collected whenever available, and nucleic acids may have degraded depending on how much time had passed since the animal defecated. Further sample collection efforts could make a more explicit effort to collect fresh (i.e. still moist) samples only.

The two viruses that were isolated from rock dove swab samples in January to March 2021 were classified as class II APMV-1 viruses of sub-genotype VI.2.1.1.1. Whether the detection of this sub-genotype reflects a dominance of this virus within urban rock doves is highly speculative, since no other genetic data for APMV-1 in New York is available. However, it is notable that there was a 2.4% percent divergence between the F gene coding sequences of these two isolates, both isolated from rock doves rescued in Manhattan, NY. This suggests that the two infections were not related but acquired independently by the two birds. It also suggests the presence of significant diversity in established and circulating strains of NDV in New York City pigeons, as well as frequent, common infection in wild birds.

Although both of our isolates contained a polybasic cleavage site in their F protein **(Figure 2)**, a molecular indicator of virulence, it is notable that neurologic clinical signs were observed in only 50% of the birds that tested positive for APMV-1 on PCR. Possible explanations include the presence of variably neurotropic strains of NDV infecting the pigeon population, or variable immune response and severity of clinical signs in a bird population adapted to this virus. Either hypothesis indicates frequent infection of wild rock doves by NDV strains in New York City.

This study found bird swab samples that tested positive for APMV-1 across Manhattan, Brooklyn, and Queens **(Figure 1)**, suggesting widespread geographic distribution of NDV across the New York City metro area. Phylogenetic characterization of our two APMV-1 isolates suggests that they are closely related to viruses isolated between 2012 and 2014 in neighboring Atlantic Flyway states: MD, PA, MI and MA. As expected, our isolates were less closely related to PPMV-1 isolates from Midwestern states along the Central Flyway, such as Kansas and Texas. Highly adapted to urban areas with increased, dense human activity, rock doves are not migratory, but have the ability to fly great distances of up to 1800 kilometers if displaced from their homes (28, 29). As a major stop along a migratory flyway, New York City’s urban and natural areas alike provide many opportunities for wild birds to forage together and otherwise interact. Therefore, the possibility that the sub-genotypes described here originated outside of New York City and were introduced through migratory, infected birds shedding those viruses cannot be excluded.

These findings indicate a need for more intensive surveillance in this region, specifically for APMV-1 but also for avian diseases generally. If circulating strains of NDV frequently infect NYC pigeons, and if the most closely related documented sequences in the Northeast date back to 2013 (**Figure 4**), future study of this population can contribute to our understanding of NDV’s evolution and genetic diversity. NDV surveillance can also inform routine husbandry and vaccination programs for backyard and commercial poultry in NY state. Finally, New York City live poultry markets and licensed small slaughterhouses should be encouraged to follow NYC Department of Health and NY State Department of Agriculture public health guidance at all times, especially when disease is detected in their flocks.

It is important to note that APMV-1 infections in humans are rare and typically cause no symptoms in humans. If symptoms occur, they are mild and self-limited influenza-like symptoms or conjunctivitis that clear up quickly with no treatment required. In fact, the lentogenic NDV vaccine strain LaSota has been proven to be safe in humans, and is used as an oncolytic agent and a vaccine vector (30–34). Although no evidence to support human-to-human transmission exists, the potential for human-to-bird transmission cannot be excluded, especially in immunocompromised individuals or those working closely with live poultry. Employers and occupational health professionals at the human-animal interface should be educated on biosafety as well as best practices around zoonotic disease transmission specific to their area of work. In the future, close monitoring of wild birds in urban settings will be essential to protecting animal and ecological safety, promoting the healthy development of poultry farming, and preventing and controlling any large-scale outbreaks of Newcastle disease virus. Community Scientists such as the New York City Virus Hunters are untapped drivers to fill in knowledge gaps for research advancement in infectious disease surveillance and communication of public health practices to the public

## Acknowledgements

This research was supported by FluLab (https://theflulab.org/about). We thank Brady Simmons of New York City Parks and Recreation for her support in obtaining sampling permits. We thank Maya Sasao and Clara Arndtsen of the Wild Bird Fund for assistance in handling wild birds. We also gratefully acknowledge Meagan McMahon, Philip Meade, and Irene Hoxie for helpful discussions around study design and data analysis.

## Author Contributions

I.F., S.B., T.B, D.D., J.G., J.G., P.K.A, F.K, R.M. and C.M. contributed to overall surveillance design. I.F., S.B., T.B., D.D., J.G., J.G. and C.M. designed and performed field data collection and laboratory analysis for the surveillance samples. I.F., C.M., E.R. and F.K. analyzed the data and drafted the manuscript. R.A. handled reporting to the USDA and BSL3 sample management. All authors reviewed the manuscript and provided critical feedback.

---

**Supplemental Table S1. Estimates of evolutionary divergence between isolates and closely related NDV F protein sequences**. The nucleotide distances between sequences are shown. Analyses were conducted using the Maximum Composite Likelihood model. This analysis involved 76 nucleotide sequences. Codon positions included were 1^st^+2^nd^+3^rd^+Noncoding. All ambiguous positions were removed for each sequence pair (pairwise deletion option). There were a total of 1656 positions.

